# PandoraRLO: DQN and Graph convolution based method for optimized ligand pose

**DOI:** 10.1101/2023.03.12.532268

**Authors:** Justin Jose, Ujjaini Alam, Divye Singh, Nidhi Jatana, Pooja Arora

## Abstract

Predicting how proteins interact with small molecules is a complex and challenging task in the field of drug discovery. Two important aspects in this are shape complementarity and inter molecular interactions which are highly driven by the binding site and the ultimate pose of the ligand in which it interacts with the protein. Various state of the art methods exist which provide a range of ligand poses that are potentially a good fit for a given specific receptor, these are usually compute intensive and expensive. In this study, we have designed a method that provides a single optimized ligand pose for a specific receptor. The method is based on reinforcement learning where when exposed to a diverse protein ligand data set the agent is able to learn the underlying complex biochemistry of the protein ligand pair and provide an optimized pair. As a first study on usage of reinforcement learning for optimized ligand pose, the PandoraRLO model is able to predict pose within a range of 0.5Å to 4Å for a large number of test complexes. This indicates the potential of reinforcement learning in uncovering the inherent patterns of protein-ligand pair in 3D space.

## Introduction

Predicting the binding of proteins and small molecules is a difficult task in drug design and can greatly impact the cost and time it takes to develop a new drug [1]. X-ray crystal structures provide important information about the binding site and the potential positions, or poses, of the ligand. However, even when the binding site is known, there may be multiple poses that a ligand can take. The binding strength, or affinity, of a ligand depends on its pose, so determining the most likely poses is crucial for protein ligand interaction [2].

Computational methods, such as those using deep learning, can greatly narrow down the search for potential drugs, but they must also be able to quickly scan through a wide range of biological and chemical possibilities [3].

Protein-ligand interactions are often highly specific and dependent on the precise shape and chemical properties of the binding site and the ligand, which can be difficult to predict [4]. Additionally, many proteins have large internal flexibility, meaning that they can adopt multiple conformations. The binding pose of a ligand can depend on the specific conformation of the protein at the time of binding, which can be difficult to predict. The binding process can also involve different types of chemical interactions between the atoms of the ligand and protein, such as hydrogen bonding, electrostatic interactions, and hydrophobic interactions, which can be difficult to predict and understand [5].

All these factors make the prediction of binding poses a challenging task for computational methods, requiring a combination of different approaches. Machine learning and AI driven methods have seen an increase in their usability as an aid to the existing computational methods, or as the primary drivers for predicting binding affinity and ligand conformations [6].

As an aid to existing docking algorithms, the AI driven methods provide enhanced scoring functions through the use of knowledge base, and extensive features other than the empirical methods [7]. AI driven scoring functions like the RF-score, NNScore and CScore can train on a set of active and inactive ligands to generate scoring function with high accuracy in distinguishing between known ligands. One of the limitation of classical machine learning based approaches is the need to represent molecules with fixed-length feature vectors. In contrast deep learning based approaches like TopologyNet [8], Pafnucy [9] and the CNN model from [10] outperform other methods in the prediction of protein-ligand binding affinities. This is due to their ability to automatically extract features directly from the two or three dimensional structures of the protein-ligand complex. Apart from scoring functions, AI approaches have also found use in overall virtual screening [11, 12, 13].

AI driven approaches greatly benefit from feature vectors of defined length. Due to their ability to provide learnable feature vectors (fingerprints) of defined length for molecules of varying sizes [14, 15, 16, 17, 18], graph based methods are finding increased use in the AI driven approaches, primarily for representation. These methods represent the three dimensional molecular structure as a 2D graph. For example, a graph embedded representation of protein-ligand pair using cascade convolution calculation method has been proposed in [19]. The use of graph based AI methods have also grown in ligand pose prediction and ranking. [20] describes the use of independently processed ligand graph and protein surface mesh representation for the prediction of binding conformation. A flexible docking mechanism using graph convolution network on atomic and *Cα* residue graph is presented in [21]. The atomic graph with basic atom features enables re-ranking poses and the *Cα* network with synthetic edges generates the distance matrix for rank prediction. [22] and [23] present a method for the proteinligand binding problem using a kNN graph representation with SE(3)-equivariant geometric deep learning model.

Reinforcement learning (RL), which is a well known technique in optimization, has in recent times found wide spread usage in the protein structure world, predominantly as an assistive technique in generative drug design [24]. Deep reinforcement learning which is a combination of RL and artificial neural network has been used to generate targeted chemical libraries of novel compounds optimized for single or multiple features[25]. RL has also been used for denovo compound generation for SARS-COV2 M protease[26]. In [27] the authors propose the use of RL as a one shot optimization approach for the protein-ligand docking problem along a single spatial coordinate. In contrast, [28] proposes a trainable 3D CNN-based RL algorithm for the rigid protein-ligand docking problem. In [29] the authors have explored the used voxelbased representation, with a single-atom ligand (copper ion). Recently, [30] used graph representation for protein and ligand with a Graph Neural Network (GNN) to obtain a learned model for addressing the protein-ligand docking problem using RL.

In this work, application of RL to predict optimized ligand pose for protein-ligand complexes is explored. The Pandora-Reinforcement Learning Optimizer (PandoraRLO) model is described in the following sections: section 2 provides the dataset and algorithms used for the model, section 3 reports the results and analysis for the same, and section 4 summarizes the conclusions drawn from the exploration of the PandoraRLO model.

## Materials and methods

This section elaborates the dataset selection, the pre-processing and preparation process along with the network design and the training and testing strategies used in PandoraRLO. Figure 1 elaborates on the overall training workflow of PandoraRLO

**Fig. 1.**
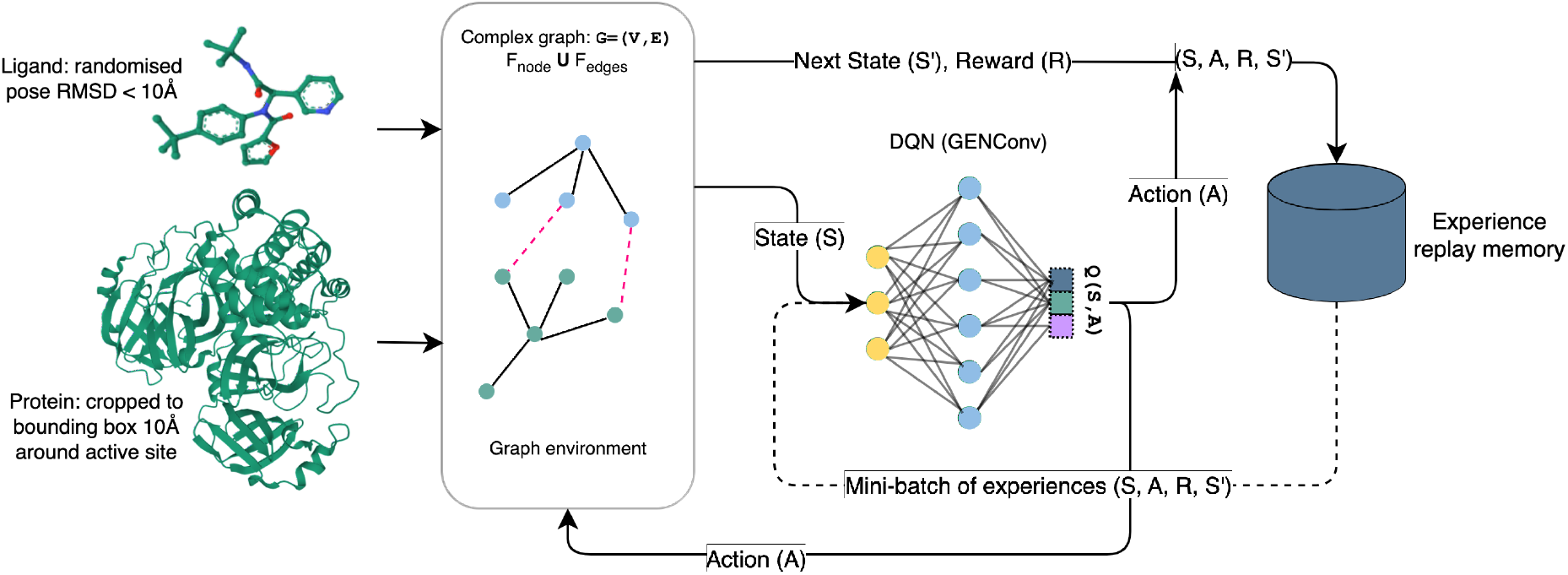
PandoraRLO training workflow: Protein ligand preparation, graph representation and DQN training

### Dataset

The dataset for PandoraRLO consists of complexes curated from the PDBBind database v2020 [31] and consists of 10, 019 protein-ligand complexes. For training and testing, PandoraRLO uses the cropped protein around the bound ligand pose, as this region contains the most relevant interactions. Also, processing the full protein-ligand complex to obtain the optimized pose would substantially deteriorate time efficiency due to a larger representational footprint. The complex is therefore cropped into a 10Å × 10Å × 10Å cube around the bound ligand centroid. The cropped complexes are protonated and charged using open babel [32].

As the final step in data pre-processing, we group the protein-ligand complexes based on the protein family information from the annotations section in PDB [33]. The training dataset comprises of 8300 complexes sampled across the protein families, and a disjoint set of 1719 complexes forms the testing dataset. The partitioning along protein families ensures a fair representation and diversity in the protein-ligand complexes presented for training and testing.

### Data Representation

PandoraRLO uses graph convolution networks for processing the protein-ligand complex, which expects the data to be represented graphically. To achieve this we convert the cropped data into a heterogeneous graph 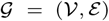, where a node represents an atom - in case of ligands, and a residue (*Cα*) in case of proteins. The edges consist of covalent bonds already present in the structure file - obtained using open babel [32] for both protein and ligand. Additional edges are added between protein residues and ligand atoms to represent their spatial proximity. This includes all edges between a protein residue and ligand atom within a distance of 4Å. The graph representation of the protein-ligand complex is built with the help of NetworkX[34].

#### Data Features

The node feature set varies based on the type of molecule. The atom node features include the atom type encoding, spatial coordinates, Van-der Waal radius, hybridization and SMARTS patterns. The residue node feature consists of residue label encoding and the atomic features of *Cα*. The atom node features are further zero padded to match the residue node feature length.

The edge attribute includes the distance encoded with Gaussian basis function with 15 different variances as described in [35].

During the course of the training, the node features and covalent bond features remain static, whereas the ligand coordinates, and the protein-ligand edge distances change depending on the actions taken by the RL agent.

### Network Design

The learning network of PandoraRLO takes graph representation of protein-ligand complex as input and outputs action which updates ligand pose. One of the major challenge in modelling the ligand pose prediction is that the actions taken to move between different ligand poses are continuous in nature. PandoraRLO uses Double Deep Q-Network (DQN) [36, 37], an off policy reinforcement learning algorithm that uses neural networks to approximate the action-value function in a *Q*-Learning framework, and demonstrates stable learning. To simplify search space for optimal ligand pose and reduce training complexity, PandoraRLO considers only translation. Since the DQN outputs action in a discrete space, the ligand pose update is discretized for a step size of 0.1Å for translations in the *x, y* and *z* axis.

PandoraRLO uses graph convolution to extract useful information from the input protein-ligand graph. The Q-learning network consists of four DeepGCN layers (DeepGCNLayer) [38] which implements skip connections with the pre-activation residual connection. Each DeepGCNLayer also uses a layer normalization, a relu activation and a GENeralized Graph Convolution (GENConv) layer [39]. The GENConv uses two MLP layers with layer normalization, dynamically learning the initial inverse temperature for softmax aggregation. GENConv allows the graph convolution to include edge features along with the node features through message passing, and captures the edge information by summing the edge features with the node features of member nodes of that edge.

### Deep Q-Network Optimization

In the DQN algorithm, the optimal *Q*-value, and the associated loss are given by:

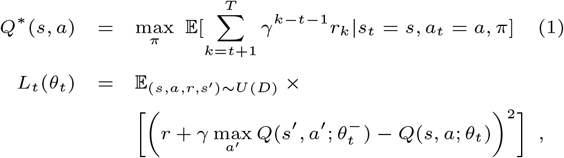

where *Q*\*(*s,a*) is the optimal state-action value which is expectation of sum of rewards *r_t_*, discounted by *γ* at each time-step *t*, maximised over policy *π*. The reward at each timestep can be realized by making an observation *s* and taking an action a, using a behaviour policy *π* = *P*(*a*|*s*). The *Q*-learning updates are applied using samples (*s, a, r, s′*), drawn episodes randomly and uniformly from the stored sample pool *D* as shown in Figure 1. The loss at the *t*-th time-step, *L_t_*(*θ_t_*), is dependent on the parameters of the *Q*-network, *θ_t_*, and the network parameters used to compute the target, 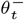, at the *t*-th timestep.

#### Reward Function

The reinforcement learning mechanism behaves as an optimization function by maximizing the overall reward. The rewards obtained at each step of the training process enables the reinforcement learning algorithm to adopt a policy which helps the ligand approach the crystallographic bound pose. Several possibilities for the reward function were considered, and the best results were obtained from a function of the RMSE between the current ligand pose and the bound pose. A relative RMSE reward function as described in equation 2 is found to perform well in predicting the optimised pose, where *RMSE*_*t*–1_ – *RMSE_t_* represents the delta change in the ligand pose RMSE post action.

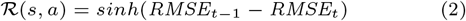

The relative RMSE based reward function enables the RL agent to get feedback based on the delta change in RMSE that every action brings instead of the RMSE itself. Further, the sine hyperbolic function helps to exponentially exaggerate the change. Thus, over several iterations, the agent learns to identify actions which would bring the ligand closer to the bound pose for a variety of complexes at different random poses.

#### Training Strategy

The protein-ligand docking problem is different from any other RL problem where either the path or the final destination is known. The differences between each complex, as highlighted in [30] makes it difficult for the RL agent to learn and generate a reusable model due to the diversity in the protein-ligand complexes, especially for unseen ones.

The conventional approach to training the reinforcement learning setup for binding pose prediction problem would be to take graphical representation of a protein-ligand complex with a random starting pose and pass it through the learning network to get action. Using this action, conformation of the selected protein-ligand complex is updated. This is repeated for either a pre-defined number of steps or until the protein-ligand complex converges to a conformation. In next episode, a new complex is taken and above process is repeated. Another method is to take more than one complex through the sequence of steps per episode. The shortcoming of these methods include:

- When the number of protein-ligand complexes increase, the learning process would have to go through large number of episodes
- Since the physico-chemical interactions change with each complex, learnings from one sequence of steps for one complex might not be directly applicable to another complex.

To overcome these shortcomings, PandoraRLO uses a learning approach where each step in any given episode, observes, processes and predicts the optimal action on distinct protein-ligand complexes. The relativistic nature of the reward function (equation 2) ensures that the scope of the reward obtained can completely be confined within each individual step. Benefit of this approach is that since in every step RL agent encounters a different complex with randomised intitial pose, it is able to explore more diverse interactions within a single episode. In this way the RL agent is able to focus on learning what makes an action better across various complexes and their interactions. Also, this approach enables the use of very large dataset in the training process.

#### Testing Strategy

For validation, the trained RL model is applied on the test dataset to predict the optimal ligand poses. The testing is done over 20 episodes per complex, with each episode predicting the optimal binding pose from a randomised initial ligand pose. The standard deviation of RMSE (*σ_RMSE_*) over a sliding window of 50 steps determines if the model has achieved convergence. A *σ_RMSE_* value less than 0.15Å is indicative of an oscillatory behaviour in the predicted actions, which suggests that the model is unable to find a better action than the ones being predicted, thus indicating model convergence for the given protein-ligand complex. In cases where the model fails to converge, the episode terminates after 400 steps.

## Results and Discussion

In order to assess the performance of the PandoraRLO model, first, the results for the training dataset have been analyzed. Figure 2 shows the loss curve for the model training. The training loss starts from a value of 4.5 and stabilizes at around 2 till 1500 episodes. At this point, for a variety of complexes in different random poses, the model is able to correctly move in the direction of the ligand bound pose ~ 96% of the time. This shows that the model has learned, to a high degree of accuracy, to move in the correct direction for the training dataset. For this convergent model, the model performance on the test dataset mentioned in Section 2.1 is examined.

**Fig. 2.**
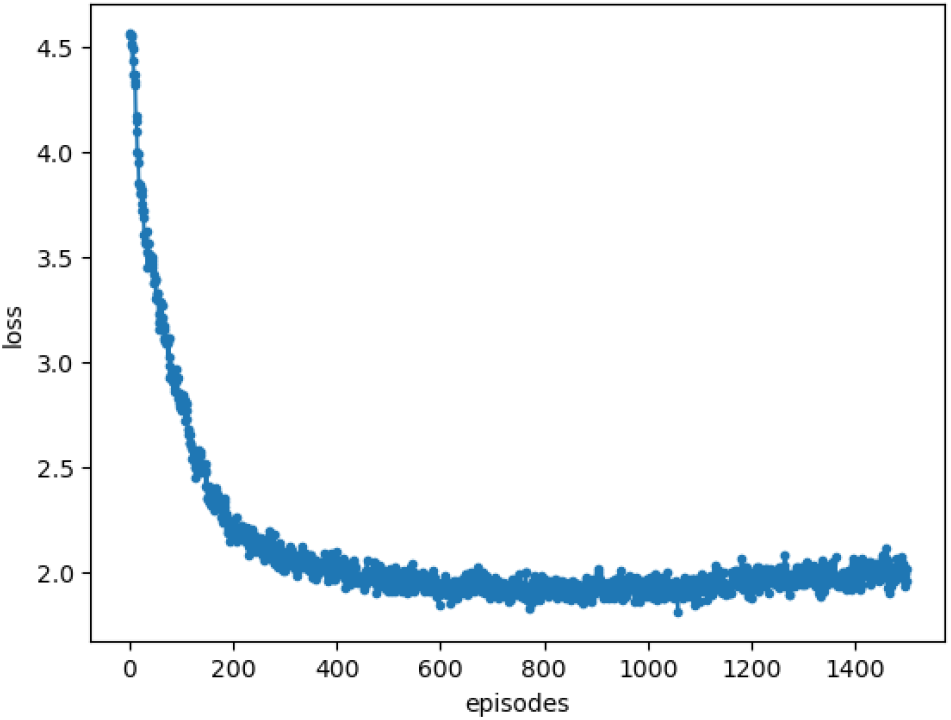
The loss curve for PandoraRLO training model.

As explained in Section 2.4.3, each test complexes starts with 20 random poses for testing. The median convergent RMSE achieved over all poses of a complex is considered to be the final RMSE attained for that complex. It is seen that the model consistently provides a median RMSE of less than or equal to 4Å for a large portion of the dataset. For the 1719 complexes considered, 28% of the data has a median RMSE of less than 2Å from the bound pose of the protein-ligand complex, while 66% of the data has a median RMSE of less than 4Å. The standard deviation for the convergent RMSE is below 1Å for 97% of the complexes which converge to below 4Å. This validates the use of median RMSE to depict the conclusive optimized pose for the different complexes. Table 1 provides a snippet of the results showing the range of final RMSE for the pose optimized by PandoraRLO. One complex has a convergent RMSE of 4.06Å, while the others have RMSE close to or less than 1Å. Three complexes from this table are also represented in the Figure 3 which shows the superimposition of the final pose trained by PandoraRLO to the bound pose in the crystal structure. We see that in all three cases, the complex approaches very close to the bound pose.

**Fig. 3.**
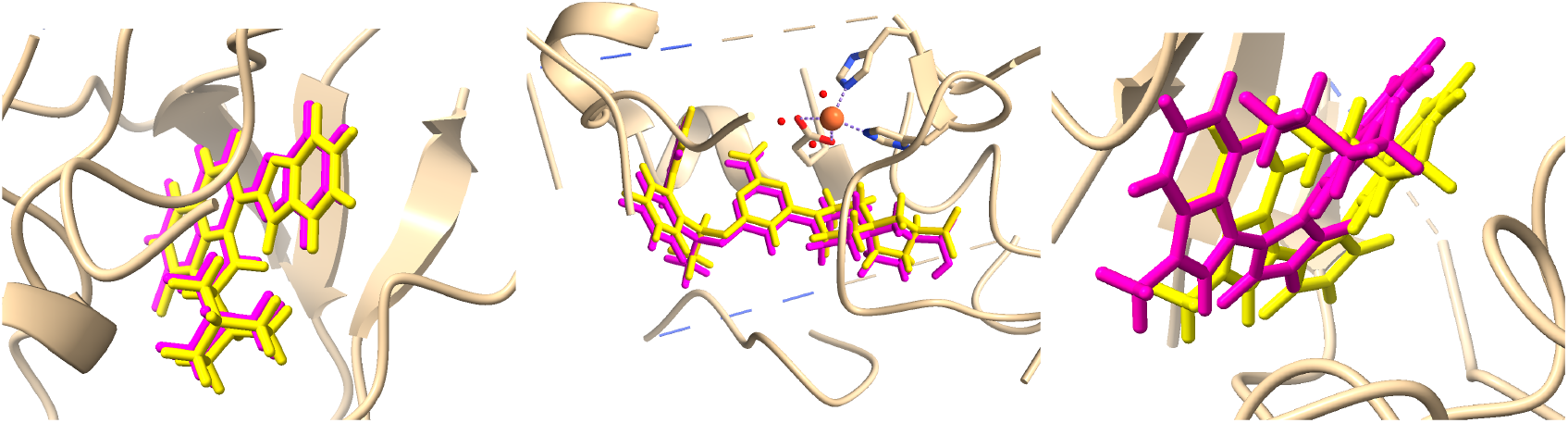
Bound (yellow) and optimized (pink) poses for the 2GDO(RMSE:1.14), 5LO1(RMSE:0.49), 1UVR(RMSE:4.04) complexes (from left to right).

**Table 1.** Model performance for sample test data. The columns represent the complex, model convergence, median RMSE (Å) and standard deviation of RMSE (Å) across 20 episodes.

| Complex ID | Convergence (%) | Median RMSE | Standard deviation |
| --- | --- | --- | --- |
| 5LO1 | 100 | 0.49 | 0.51 |
| 5NGU | 100 | 0.50 | 0.30 |
| 4DPI | 100 | 0.92 | 0.11 |
| 1BDR | 100 | 1.06 | 0.27 |
| 2GDO | 100 | 1.14 | 0.09 |
| 4MUW | 100 | 1.21 | 0.21 |
| 1UVR | 90 | 4.06 | 0.16 |

To further study the results, multiple smaller experiments were conducted to understand the learning behavior of the model. These included dividing the entire train-test data based on protein families, training the model for an individual protein family, segregating similar types of ligands and train-testing on them, and training the model on clusters formed using model features. There was no consistent correlation between the RMSE and various data subsets. It was, however, observed in the test results that proteins from over represented families in the training dataset show consistently lower RMSE at testing as compared to others, underlining the importance of greater availability of datasets from different protein families.

Next, the feature set for the trained model was analyzed. GNNExplainer [40], which is an explanation tool for graph neural network based predictions, was used to understand which features are given importance by the learning network while taking an action. For illustrative purposes the results for two complexes, 2GDO and 1UVR, are shown in Figure 4. The residues and edges identified by GNNExplainer were cross validated with the experimental data using Ligplot [41], as shown in Figure 5.

**Fig. 4.**
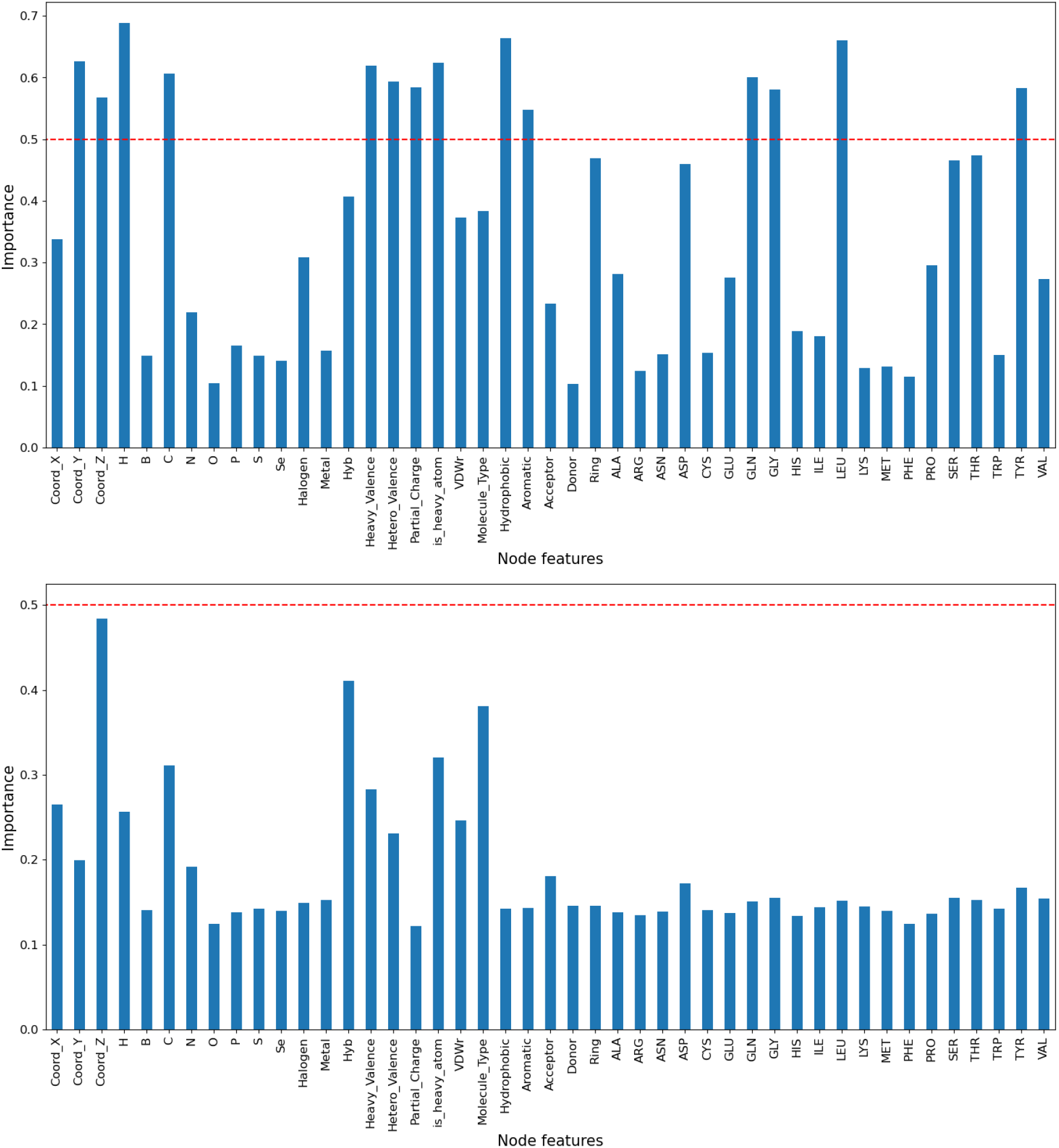
GNNExplainer feature importance at 1500 episodes for 2GDO (top) and 1UVR (bottom).

**Fig. 5.**
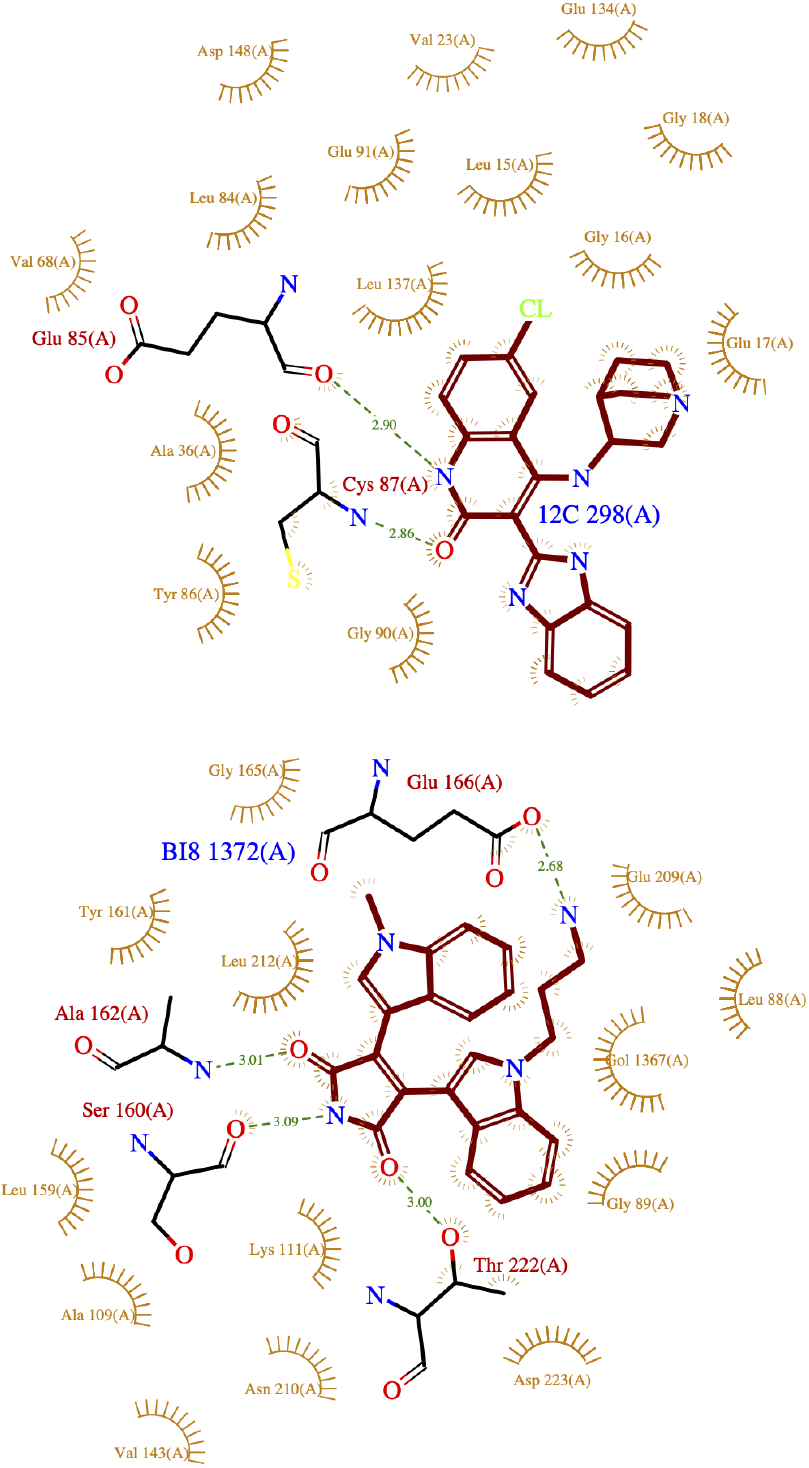
Ligplots for 2GDO (top) and 1UVR (bottom).

For 2GDO, where the final RMSE was 1.14Å, the list of features identified as important by GNNExplainer included the coordinates, H, C, heavy valence, hetero valence, partial charge, hydrophobic and aromatic bonds, as well as the residues Glutamine (GLU), Glycine (GLY), Leucine (LEU) and Tyrosine (TYR) and their inter-molecular edges. The above residues from GNNExplainer match some of the major interacting residues as seen in Ligplot. It should be noted that some interacting residues such as Alanine (ALA) and Cysteine (CYS) were not given particularly high importance. However, overall, the model appeared to have correctly given greater weightage to the relevant interacting residues.

For the other complex 1UVR, where the final RMSE was 4.06Å, no feature was given importance over 0.5, the highest importance was given to the z-coordinate and hybridization. None of the residues were considered significant (less than 0.2 importance). The interacting residues seen in the Ligplot, such as GLU, GLY, SER, THR, etc, were not given enough weightage in GNNExplainer.

The above GNNExplainer analysis indicates that for 2GDO, where the final RMSE is low, the model gave importance to most of the relevant interacting residues of the bound pose. In contrast, for 1UVR, where the final RMSE was high, the model was not successful in picking up the relevant residues.

This seems to suggest that the learning network is able perform well when it can extract meaningful interactions from the feature set. Further examination of these results may lead to a deeper understanding of the importance of the feature set for RL optimization of protein-ligand docking.

### Model evaluation against existing tools

#### Equibind

To validate the PandoraRLO model against already published research and existing tools, the model was tested against Equibind [42] which was published early last year and has claimed to show better performance than many state of the art tools. The PandoraRLO trained model was tested against the Equibind publicly available test data of 358 complexes. The results from PandoraRLO and Equibind models were compared based on the final pose of the ligand molecule using centroid separation. Out of the 358 complexes, PandoraRLO and Equibind predict a centroid separation less than 2Å for 113 and 134 complexes respectively. Similarly for complexes where the centroid separation converged to less than 4Å, PandoraRLO and Equibind predictions are 236 and 222 respectively. For a significant fraction of the data, PandoraRLO provides a centroid separation lower than or comparable to Equibind. In table 2, we provide a snapshot of the results from both models for a few complexes, for some of which PandoraRLO performs better, while for others Equibind gives better results. From the overall results, we can say that the PandoraRLO model, which is only translation based with a rigid protein-ligand pair, has demonstrated comparable performance to Equibind.

**Table 2.** Comparison on sample Equibind test data based on centroid separation

| Complex | centroid separation (in Å) |  |  |
| --- | --- | --- | --- |
|  | Starting | Equibind | PandoraRLO |
| 6NXZ | 13.920 | 3.248 | 1.022 |
| 6G5U | 15.929 | 2.204 | 1.011 |
| 6NY0 | 12.928 | 3.264 | 0.487 |
| 6I65 | 14.519 | 1.324 | 1.715 |
| 6PNM | 14.555 | 1.004 | 1.027 |

#### Autodock-Vina

The samples mentioned in the table 2 were also evaluated using Autodock-Vina [43]. Autodock-Vina docking tools were used to identify the highest ranked pose with lowest binding energy for the given protein-ligand pair. For these complexes, PandoraRLO was able to achieve an RMSE in the range of 0.5-2 Å, whereas Autodock-Vina shows an RMSE of around 3 Å. Given that PandoraRLO considers rigid protein and rigid ligand, it was observed that the pose generated by PandoraRLO were comparable to Autodock-Vina. Figure 6 and 7 illustrates the comparison of poses generated by PandoraRLO and Autodock-Vina with the bound pose.

**Fig. 6.**
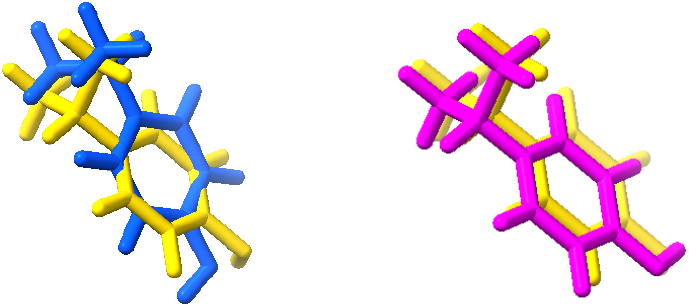
Bound pose (yellow), Optimized pose by Vina (Blue) and optimized pose by PandoraRLO(pink) for PDBid:6I65

**Fig. 7.**
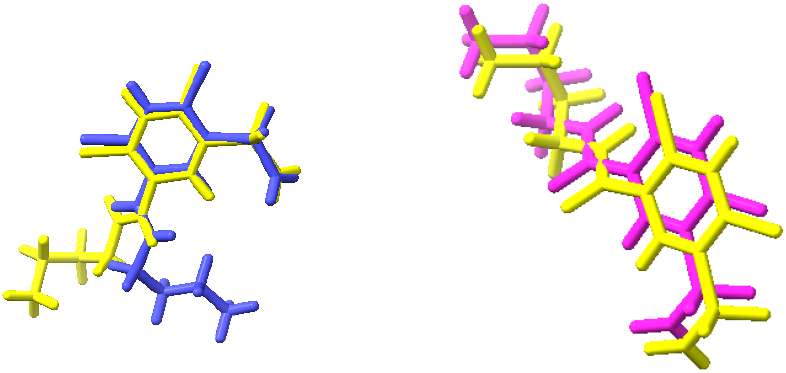
Bound pose (yellow), Optimized pose by Vina (Blue) and optimized pose by PandoraRLO(pink) for PDBid:6G5U

## Conclusion

In this work, a reinforcement learning-based model, PandoraRLO, has been proposed for obtaining optimized ligand pose. The model utilizes a graph convolution-based method for representation and an RMSE-based reward function. This model has demonstrated promising results compared to existing deep learning and state-of-the-art methods. PandoraRLO provides a convergent model trained on a dataset of 8300 complexes from various protein families. This when tested on 1719 test complexes is able to converge to an RMSE of ≤ 4Å for 66% of the test complexes. Interestingly, preliminary analysis of a small sample of the results seems to suggest that the model is able to, at least partially, choose the relevant inter-molecular interactions from the feature set. Thus this model has the potential to unveil the ability of an RL agent to learn the underlying structural and network patterns. As future work, we would like enhance current model to incorporate rotation and combine network properties with biochemical properties to increase the biological relevance of the optimized pose. This may result in the RL agent being able to understand the nuances and uniqueness of protein-ligand interactions to create computer-based predictions that are close to real-world experiments.

## Availability and reproducibility

The code and the dataset used is available at https://github.com/thoughtworks/PandoraRL

## Acknowledgements

We would like to thank Geetansh Kalra for his contribution and help during the data curation and preparation stage.

## References

1. Xing Du, Yi Li, Yuan-Ling Xia, Shi-Meng Ai, Jing Liang, Peng Sang, Xing-Lai Ji, and Shu-Qun Liu. Insights into protein–ligand interactions: mechanisms, models, and methods. International journal of molecular sciences, 17(2):144, 2016.

2. Hiraku Oshima, Suyong Re, and Yuji Sugita. Prediction of protein–ligand binding pose and affinity using the grest+ fep method. Journal of Chemical Information and Modeling, 60(11):5382–5394, 2020.

3. George Lambrinidis and Anna Tsantili-Kakoulidou. Multiobjective optimization methods in novel drug design. Expert Opinion on Drug Discovery, 16(6):647–658, 2021.

4. Yi Fu, Ji Zhao, and Zhiguo Chen. Insights into the molecular mechanisms of protein-ligand interactions by molecular docking and molecular dynamics simulation: A case of oligopeptide binding protein. Computational and mathematical methods in medicine, 2018, 2018.

5. Sangmin Seo, Jonghwan Choi, Sanghyun Park, and Jaegyoon Ahn. Binding affinity prediction for protein–ligand complex using deep attention mechanism based on intermolecular interactions. BMC bioinformatics, 22(1):1–15, 2021.

6. Zechen Wang, Liangzhen Zheng, Sheng Wang, Mingzhi Lin, Zhihao Wang, Adams Wai-Kin Kong, Yuguang Mu, Yanjie Wei, and Weifeng Li. A fully differentiable ligand pose optimization framework guided by deep learning and traditional scoring functions. arXiv preprint arXiv:2206.13345, 2022.

7. Pin Chen, Yaobin Ke, Yutong Lu, Yunfei Du, Jiahui Li, Hui Yan, Huiying Zhao, Yaoqi Zhou, and Yuedong Yang. Dligand2: an improved knowledge-based energy function for protein–ligand interactions using the distance-scaled, finite, ideal-gas reference state. Journal of cheminformatics, 11(1):1–11, 2019.

8. Zixuan Cang and Guo-Wei Wei. Topologynet: Topology based deep convolutional and multi-task neural networks for biomolecular property predictions. PLOS Computational Biology, 13(7):e1005690, 2017.

9. Marta M Stepniewska-Dziubinska, Piotr Zielenkiewicz, and Pawel Siedlecki. Development and evaluation of a deep learning model for protein–ligand binding affinity prediction. Bioinformatics, 34(21):3666–3674, 05 2018.

10. Matthew Ragoza, Joshua Hochuli, Elisa Idrobo, Jocelyn Sunseri, and David Ryan Koes. Protein-ligand scoring with convolutional neural networks. Journal of Chemical Information and Modeling, 57(4):942–957, 2017.

11. Liwei Li, May Khanna, Inha Jo, Fang Wang, Nicole M. Ashpole, Andy Hudmon, and Samy O. Meroueh. Target-specific support vector machine scoring in structurebased virtual screening: Computational validation, in vitro testing in kinases, and effects on lung cancer cell proliferation. Journal of Chemical Information and Modeling, 51(4):755–759, 2011.

12. David Xu and Samy O. Meroueh. Effect of binding pose and modeled structures on svmgen and glidescore enrichment of chemical libraries. Journal of Chemical Information and Modeling, 56(6):1139–1151, 2016.

13. Nan Li, Richard I. Ainsworth, Meixin Wu, Bo Ding, and Wei Wang. Miec-svm: Automated pipeline for protein peptide/ligand interaction prediction. Bioinformatics, 32(6):940–942, 2015.

14. A. Ahmed, B. Mam, and R. Sowdhamini. DEELIG: A Deep Learning Approach to Predict Protein-Ligand Binding Affinity. Bioinform Biol Insights, 15:2020–09, 2021.

15. David Duvenaud, Dougal Maclaurin, Jorge Aguilera-Iparraguirre, Rafael Gómez-Bombarelli, Timothy Hirzel, Alán Aspuru-Guzik, and Ryan P. Adams. Convolutional networks on graphs for learning molecular fingerprints. In Proceedings of the 28th International Conference on Neural Information Processing Systems - Volume 2, NIPS’15, page 2224–2232, Cambridge, MA, USA, 2015. MIT Press.

16. Han Altae-Tran, Bharath Ramsundar, Aneesh S Pappu, and Vijay Pande. Low data drug discovery with one-shot learning. ACS central science, 3(4):283–293, 2017.

17. Alex M Fout. Protein interface prediction using graph convolutional networks. PhD thesis, Colorado State University, 2017.

18. Wen Torng and Russ B Altman. Graph convolutional neural networks for predicting drug-target interactions. Journal of chemical information and modeling, 59(10):4131–4149, 2019.

19. Huimin Shen, Youzhi Zhang, Chunhou Zheng, Bing Wang, and Peng Chen. A cascade graph convolutional network for predicting protein–ligand binding affinity. International journal of molecular sciences, 22(8):4023, 2021.

20. Oscar Méndez-Lucio, Mazen Ahmad, Ehecatl Antonio del Rio-Chanona, and Jörg Kurt Wegner. A geometric deep learning approach to predict binding conformations of bioactive molecules. Nature Machine Intelligence, 3(12):1033–1039, 2021.

21. Amr H. Mahmoud, Jonas F. Lill, and Markus A. Lill. Graph-convolution neural network-based flexible docking utilizing coarse-grained distance matrix. CoRR, 2020.

22. Octavian-Eugen Ganea, Xinyuan Huang, Charlotte Bunne, Yatao Bian, Regina Barzilay, Tommi S. Jaakkola, and Andreas Krause. Independent se(3)-equivariant models for end-to-end rigid protein docking, 2021.

23. Hannes Stärk, Octavian-Eugen Ganea, Lagnajit Pattanaik, Regina Barzilay, and Tommi Jaakkola. Equibind: Geometric deep learning for drug binding structure prediction, 2022.

24. Xin Yang, Yifei Wang, Ryan Byrne, Gisbert Schneider, and Shengyong Yang. Concepts of artificial intelligence for computer-assisted drug discovery. Chemical reviews, 119(18):10520–10594, 2019.

25. Mariya Popova, Olexandr Isayev, and Alexander Tropsha. Deep reinforcement learning for de novo drug design. Science advances, 4(7):eaap7885, 2018.

26. Alberga Domenico, Gambacorta Nicola, Trisciuzzi Daniela, Ciriaco Fulvio, Amoroso Nicola, and Nicolotti Orazio. De novo drug design of targeted chemical libraries based on artificial intelligence and pair-based multiobjective optimization. Journal of Chemical Information and Modeling, 60(10):4582–4593, 2020.

27. Antonio Serrano, Baldomero Imbernón, Horacio Pérez-Sánchez, José M. Cecilia, Andrés Bueno-Crespo, and José L. Abellán. Qn-docking: An innovative molecular docking methodology based on q-networks. Applied Soft Computing, 96:106678, 2020.

28. Justin Jose, Kritika Gupta, Ujjaini Alam, Nidhi Jatana, and Pooja Arora. Reinforcement learning based approach for ligand pose prediction, 2021.

29. Chenran Wang, Yang Chen, Yuan Zhang, Keqiao Li, Menghan Lin, Feng Pan, Wei Wu, and Jinfeng Zhang. A reinforcement learning approach for protein–ligand binding pose prediction. BMC Bioinformatics, 23(1):368, Sep 2022.

30. Justin Jose, Ujjaini Alam, Divye Singh, Nidhi Jatana, and Pooja Arora. PandoraRL: DQN and Graph Convolution based ligand pose learning for SARS-COV1 Mprotease. In 2022 IEEE International Conference on Bioinformatics and Biomedicine (BIBM), pages 380–385, 2022.

31. Renxiao Wang, Xueliang Fang, Yipin Lu, Chao-Yie Yang, and Shaomeng Wang. The pdbbind database: methodologies and updates. Journal of medicinal chemistry, 48(12):4111–4119, 2005.

32. Noel M O’Boyle, Michael Banck, Craig A James, Chris Morley, Tim Vandermeersch, and Geoffrey R Hutchison. Open babel: an open chemical toolbox. Journal of Cheminformatics, 3(1):33, 2011.

33. Stephen K Burley, Helen M Berman, Gerard J Kleywegt, John L Markley, Haruki Nakamura, and Sameer Velankar. Protein data bank (pdb): the single global macromolecular structure archive. Protein Crystallography, pages 627–641, 2017.

34. Aric A. Hagberg, Daniel A. Schult, and Pieter J. Swart. Exploring network structure, dynamics, and function using networkx. In Gaël Varoquaux, Travis Vaught, and Jarrod Millman, editors, Proceedings of the 7th Python in Science Conference, pages 11 – 15, Pasadena, CA USA, 2008.

35. John Jumper, Richard Evans, Alexander Pritzel, Tim Green, Michael Figurnov, Olaf Ronneberger, Kathryn Tunyasuvunakool, Russ Bates, Augustin Žídek, Anna Potapenko, Alex Bridgland, Clemens Meyer, Simon A. A. Kohl, Andrew J. Ballard, Andrew Cowie, Bernardino Romera-Paredes, Stanislav Nikolov, Rishub Jain, Jonas Adler, Trevor Back, Stig Petersen, David Reiman, Ellen Clancy, Michal Zielinski, Martin Steinegger, Michalina Pacholska, Tamas Berghammer, Sebastian Bodenstein, David Silver, Oriol Vinyals, Andrew W. Senior, Koray Kavukcuoglu, Pushmeet Kohli, and Demis Hassabis. Highly accurate protein structure prediction with alphafold. Nature, 596(7873):583–589, 2021.

36. Volodymyr Mnih, Koray Kavukcuoglu, David Silver, Alex Graves, Ioannis Antonoglou, Daan Wierstra, and Martin A. Riedmiller. Playing atari with deep reinforcement learning, 2013.

37. Volodymyr Mnih, Koray Kavukcuoglu, David Silver, Andrei Rusu, Joel Veness, Marc Bellemare, Alex Graves, Martin Riedmiller, Andreas Fidjeland, Georg Ostrovski, Stig Petersen, Charles Beattie, Amir Sadik, Ioannis Antonoglou, Helen King, Dharshan Kumaran, Daan Wierstra, Shane Legg, and Demis Hassabis. Human-level control through deep reinforcement learning. Nature, 518:529–33, 02 2015.

38. Guohao Li, Matthias Müller, Ali Thabet, and Bernard Ghanem. Deepgcns: Can gcns go as deep as cnns? In The IEEE International Conference on Computer Vision (ICCV), 2019.

39. Guohao Li, Chenxin Xiong, Ali Thabet, and Bernard Ghanem. Deepergcn: All you need to train deeper gcns, 2020.

40. Rex Ying, Dylan Bourgeois, Jiaxuan You, Marinka Zitnik, and Jure Leskovec. Gnnexplainer: Generating explanations for graph neural networks, 2019.

41. Andrew C Wallace, Roman A Laskowski, and Janet M Thornton. Ligplot: a program to generate schematic diagrams of protein-ligand interactions. Protein engineering, design and selection, 8(2):127–134, 1995.

42. Hannes Stärk, Octavian Ganea, Lagnajit Pattanaik, Regina Barzilay, and Tommi Jaakkola. Equibind: Geometric deep learning for drug binding structure prediction. In International Conference on Machine Learning, pages 20503–20521. PMLR, 2022.

43. Oleg Trott and Arthur J Olson. Autodock vina: improving the speed and accuracy of docking with a new scoring function, efficient optimization, and multithreading. Journal of computational chemistry, 31(2):455–461, 2010.

